# smDeepFLUOR: Single-Molecule Deep Learning Fluorescence Classification

**DOI:** 10.1101/2025.07.09.663809

**Authors:** Jinseob Lee, Byungju Kim, Gayun Bu, Muhammad Tehseen, Vlad-Stefan Raducanu, Samir M Hamdan, Jong-Bong Lee

**Affiliations:** Devision of Interdisciplinary Bioscience & Bioengineering, Pohang University of Science & Technology (POSTECH), Pohang 37673, Republic of Korea; Department of Physics, Pohang University of Science & Technology (POSTECH), Pohang 37673, Republic of Korea; Bioscience Program, Division of Biological and Environmental Sciences and Engineering, King Abdullah University of Science and Technology, Thuwal 23955, Saudi Arabia

## Abstract

Fluorescence intensity variation has long served as a primary readout for monitoring biological events. However, single-fluorophore signals arising from distinct molecular events often exhibit similar intensity profiles, making further classification challenging using conventional methods. In this study, we introduce smDeepFLUOR, a deep learning–based framework that resolves seemingly homogeneous spatiotemporal fluorescence signals by uncovering latent features imperceptible to conventional analyses. By leveraging a three-dimensional convolutional neural network trained on image sequences captured over 7 × 7 × 10 voxel windows, smDeepFLUOR reliably distinguishes specific from nonspecific protein binding, even across different experimental days, with an accuracy of up to 97%. Remarkably, smDeepFLUOR also captures real-time DNA synthesis kinetics by identifying subtle changes in the spatial distance between the fluorophore and the 3’ end of nascent DNA, a feature undetectable by traditional methods. These classifications were achieved without incorporating explicit physical rules or engineered features, implying the presence of intrinsic, previously unrecognized differences in emission patterns. This approach significantly extends the analytical capabilities of single-molecule fluorescence imaging and opens new avenues for minimally labeled and label-free protein activities.

## Introduction

Pixelated single-molecule fluorescence signals can offer unparalleled insights into the dynamics of individual biomolecules by analyzing their intensity profiles^1^. These signals can emerge from interactions between fluorophores and surrounding environments, leading to abrupt shifts in fluorescence signals via mechanisms such as protein-induced fluorescence enhancement or quenching (PIFE/PIFQ)^2^ and nucleic acid-induced enhancement (NAIFE)^3, 4^ or quenching (NAIFQ)^5^. Although it remains challenging to generalize fluorescence responses across diverse environments, single-fluorophore signals not only reflect the presence of a biomolecule but also exhibit inherent variations depending on their local surroundings^6^.

Under the premise that subtle environmental changes can induce distinct states in fluorophores, we investigated whether advanced deep-learning methods could classify these states, even when the average fluorescence intensity appears indistinguishable. Although fluorescence signals captured by an EMCCD camera exhibit Gaussian-like intensity profiles over only a few pixels, previous studies on single-molecule emission patterns have revealed that these intensity distributions contain more intricate information than intuitively expected^7, 8^. Modern deep-neural networks^9, 10, 11^, particularly convolutional neural networks (CNNs)^12^, have shown robust capabilities in extracting essential features and achieving remarkable classification accuracy. Recent advancements in applying deep-learning to fluorescence imaging data have enhanced traditional methods, enabling the differentiation of features that were previously considered indistinguishable^13, 14, 15, 16^.

In this study, we developed a novel approach called single-molecule deep-learning fluorescence classification (smDeepFLUOR), designed to classify distinct states of proteins or DNA labeled with a single fluorophore. This method uses ten frames of a two-dimensional (2D) image consisting of 7 x 7 pixels. The three-dimensional (3D) CNN architecture of smDeepFLUOR processes a 7 x 7 x 10 image of fluorescence intensity distribution, rather than relying on a single averaged intensity, thereby capturing comprehensive spatiotemporal data (Figure 1).

**Figure 1.**
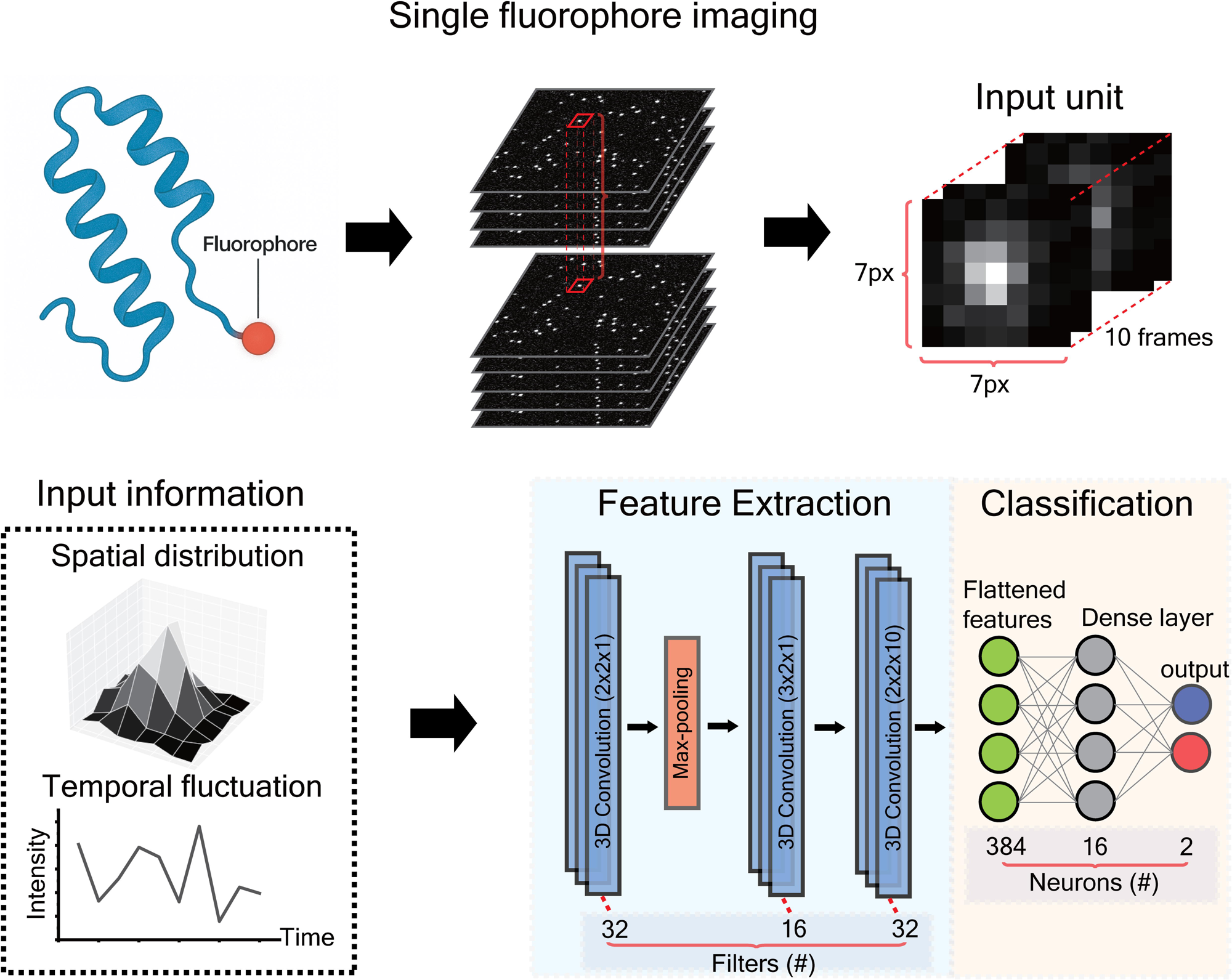
Schematic of smDeepFLUOR. Fluorophore-conjugated biomolecules are imaged using TIRF microscopy. The smDeepFLUOR estimates the spatiotemporal fluorescence signals within a 7 x 7 x 10 voxel window using a 3D convolutional neural network.

We demonstrate that smDeepFLUOR can distinguish the specific binding of poly(A)-binding protein (PABP) to poly(A) RNA tails (93% accuracy) from its nonspecific binding to the surface of an imaging chamber (97% accuracy). The trained model reliably classifies the specific (85% accuracy) and nonspecific (90% accuracy) binding of PABP across data obtained on different. Surprisingly, smDeepFLUOR can also detect subtle changes in emission patterns arising from the proximity of the 3’ end of nascent DNA to Cy3 labeled at the 5’ end, whose differences are indistinguishable using conventional methods. This smDeepFLUOR capability allows us to monitor the kinetics of DNA synthesis with a single fluorophore attached to DNA, even in the absence of discernible PIFE as DNA polymerase approaches the fluorophore. We show the capabilities of smDeepFLUOR in revealing previously unrecognized intrinsic information in single-molecule fluorescence signals, enabling a new class of fluorescence-based assays that operate beyond the sensitivity of conventional intensity-based methods.

## Results

### Classification of specific interactions by smDeepFLUOR

We first investigated whether a fluorophore directly conjugated to proteins could sense changes in the local environment arising from specific or non-specific protein interactions. To assess smDeepFLUOR to detect binding specificity, we prepared a human poly(A)-binding protein fused to a single mNeonGreen (mNG) fluorescent protein. The mNG-PABP specifically binds to the poly(A) region of a DNA/RNA partial duplex immobilized on a surface passivated with a mixture of polyethylene glycol (PEG) and PEG-Biotin (Figure 2a). Specific-binding datasets were collected by cropping mNG-PABP signals colocalized with Cy5 signals at a DNA/RNA partial duplex while mNG signals in the absence of the DNA/RNA partial duplex were collected for nonspecific mNG-PABP binding to the surface, imaged by total internal reflection fluorescence (TIRF) microscopy. Of the 40,000 image sets collected in the same sample, 80% were used to train smDeepFLUOR. Testing with the remaining 20% showed that smDeepFLUOR achieved an identification accuracy of approximately 97% and 93% for classifying nonspecific surface-bound and specific poly(A)-bound mNG-PABP, respectively (Figure 2a). Notably, despite indistinguishable pixelated image sequences and average fluorescence intensities between the two classes, smDeepFLUOR successfully distinguished them, revealing the presence of subtle yet discriminative spatiotemporal features.

**Figure 2.**
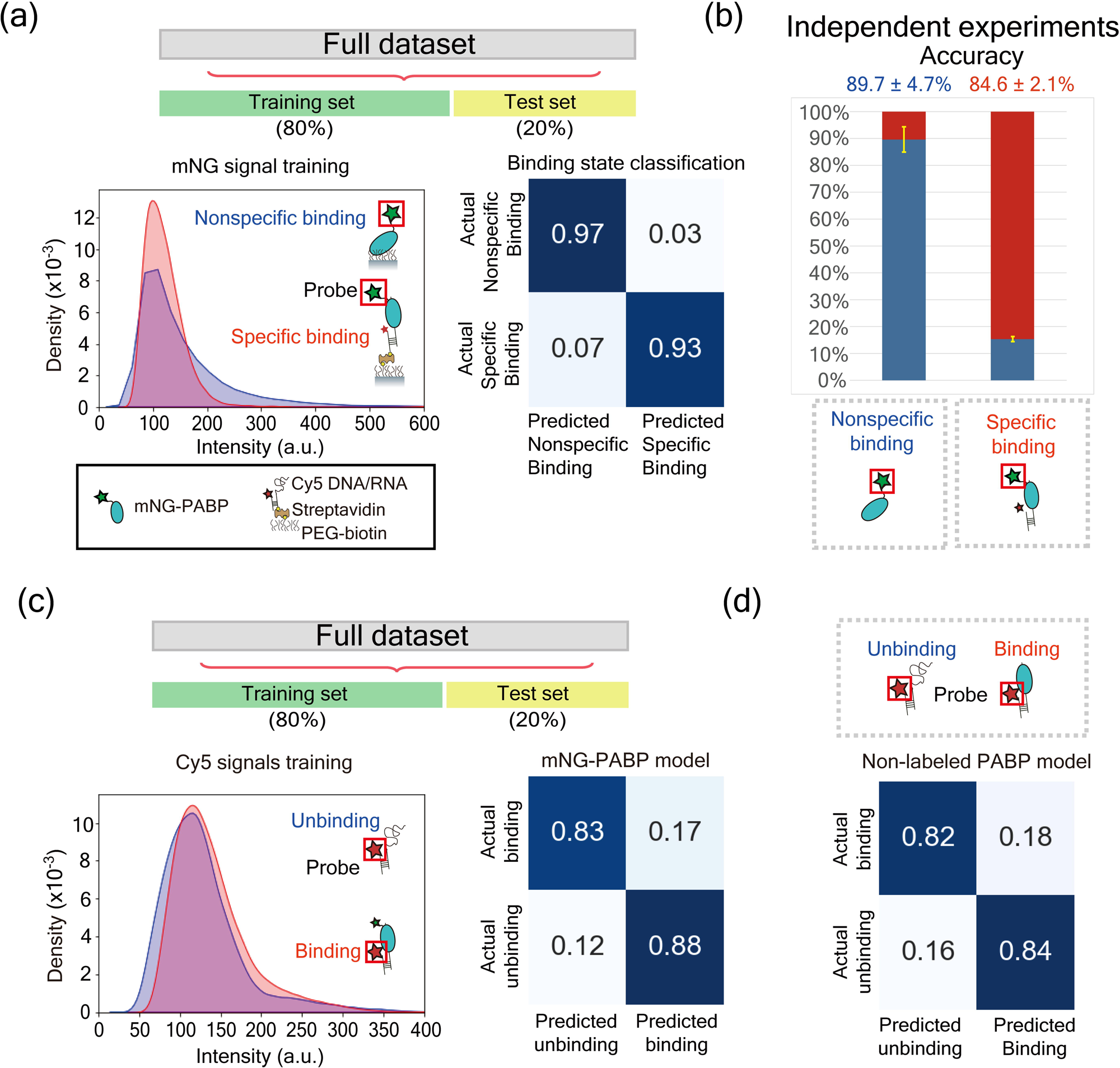
Identification of specific interactions between poly(A) tail and PABP. (a) Fluorescence signals from mNG-PABP were used for training (green star in red box). The intensity of mNG-PABP did not significantly differ between specific and nonspecific binding (histogram). The dataset was split into 80% training and 20% testing sets. The smDeepFLUOR trained on the full dataset achieved 97% accuracy for identifying nonspecific surface binding and 93% accuracy for identifying actual specific binding to the poly(A) tail. (b) The smDeepFLUOR achieved 90% accuracy for nonspecific binding and 85% accuracy for specific binding on unseen data from independent experiments conducted on different days. (c) Experimental design for classifying protein binding using mNG-PABP. Fluorescence signals from Cy5-labeled DNA/RNA partial duplex were used for training (red star in red box). Despite similar Cy5 intensities between protein-bound and unbound states (histogram), the model achieved 83% accuracy for the bound state and 88% for the unbound state. (d) Fluorescence signals from Cy5-DNA/RNA partial duplex were used for training. The model predicted the binding state with 82% accuracy and the unbinding state with 84% accuracy.

Furthermore, to evaluate whether smDeepFLUOR could reliably analyze images from independent experiments that share the common feature of being the same protein but have entirely different background characteristics^17^, we tested its from Figure 2b, smDeepFLUOR achieved an identification accuracy of 89.7 ± 4.7% performance on datasets collected on different days. When trained on the dataset from Figure 2b, smDeepFLUOR achieved an identification accuracy of 89.7 ± 4.7% (mean ± S.D.) for nonspecific binding to the surface and 84.6 ± 2.1% for specific binging to poly(A) in images obtained from these separate experiments (Figure 2b). These results suggest that even when trained on data from a limited number of experiments, smDeepFLUOR demonstrated accuracy levels comparable to those achieved with the full dataset, highlighting its robustness. Notably, smDeepFLUOR can maintain consistent performance in independent experiments.

To address the challenge of recognizing protein binding when changes in fluorescence intensity fall outside the detectable range of PIFE or PIFQ, we first trained smDeepFLUOR using mNG-PABP binding events with a single Cy5 dye attached to a DNA/RNA partial duplex (Figure 2c). This time, we tested whether a fluorophore not attached to the protein could detect environmental changes induced by a nearby binding protein. For training, PABP-bound datasets were prepared by extracting Cy5 signals colocalized with mNG-PABP while Cy5 signals in the absence of mNG-PABP were used for PABP-unbound datasets (Figure 2c). Remarkably, smDeepFLUOR achieved an identification accuracy of about 86%, with 83% accuracy for unbinding events and 88% for binding events to the template, for classifying protein binding events based on Cy5 signals on the template (Figure 2c). These results indicate that smDeepFLUOR can successfully identify specific protein binding to the template, even beyond the range of PIFE and PIFQ.

### Real-time protein binding detection without the need for labeled proteins

In our previous studies, we found that PABP could occupy over 90% of Poly(A) templates in the presence of 10 nM PABP molecules, owing to its low dissociation constant (K_d_) of about 1.0 nM^18^. This high proportion of unlabeled PABP bound data enabled us to effectively train smDeepFLUOR using unlabeled PABP molecules^19^. The resulting identification accuracy for unlabeled PABP binding reached 83% (Figure 2d). When compared to smDeepFLUOR trained on mNG-PABP binding data based on colocalization with Cy5 (Figure 2c), its identification accuracy matches well those of mNG-PABP (Figure S1b). Taken together, this result suggests the potential for broad application of this approach in training with unlabeled proteins. Since labeling the protein of interest is often a significant challenge in performing fluorescence imaging including new single-molecule experiments, we anticipate that smDeepFLUOR for analyzing unlabeled proteins will facilitate the development of innovative experimental concepts.

To validate smDeepFLUOR to classify single protein binding events in real-time, we further analyzed individual trajectories by examining the colocalization of PABP and its template: mNG-PABP was introduced into an imaging chamber where Cy5-labeled DNA/RNA partial duplexes were immobilized, unbound proteins were washed away, and then both fluorescence channels were sequentially imaged (Figure 3a). The model, trained on non-labeled PABP (Figure 2d), accurately detected the timing of mNG-PABP appearance corresponding to binding to a DNA/RNA partial duplex, even when the Cy5 intensity remained unchanged. we showed that smDeepFLUOR successfully identified PABP binding states by recognizing the spatiotemporal overlap between mNG-PABP and Cy5-DNA/RNA partial duplex although it had not been trained with fluorescently labeled PABP (Figure 3c). These results demonstrate the model’s capacity to monitor real-time protein binding at the single-molecule level, without relying on significant fluorescence intensity fluctuations and labeled proteins for training.

**Figure 3.**
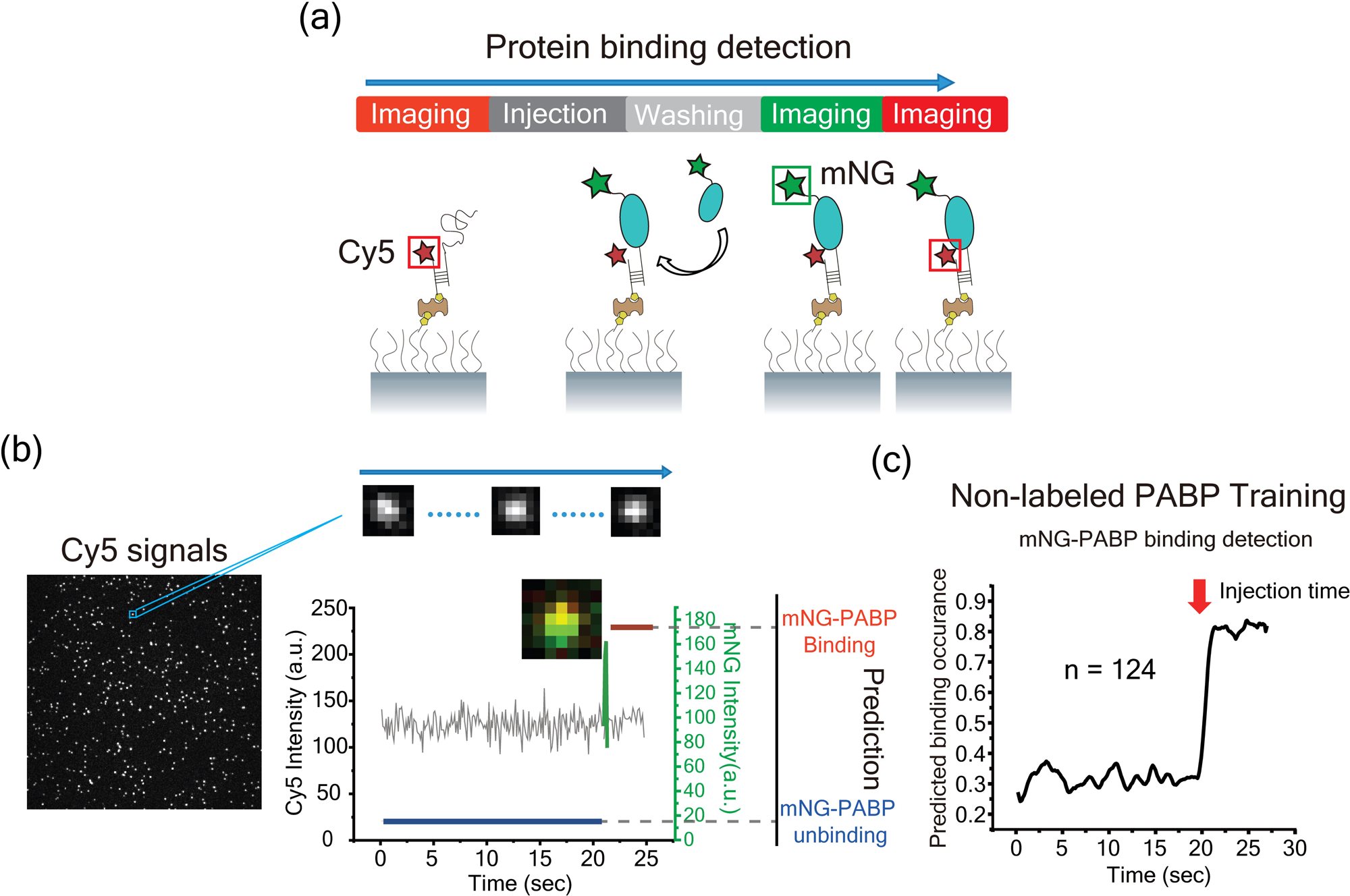
Real-time detection of PABP binding to poly(A) tail. (a) Schematic illustration of the colocalization experiment showing mNG-PABP binding to a Cy5-DNA/RNA partial duplex. (b) Representative intensity traces of a single mNG-PABP molecule (green) bound to a single Cy5-DNA/RNA duplex (gray), overlaid with model predictions of protein binding (red) and unbinding (blue) events. Using mNG-PABP as a true indicator of protein binding, we found that the moment of PABP binding could be accurately identified by predicting the Cy5 signal on the DNA/RNA partial duplex using the model. (c) Averaged prediction results of mNG-PABP traces bound to Cy5-labeled DNA/RNA partial duplexes (n = 124).

### Classification of two structurally distinct DNA constructs

To assess whether smDeepFLUOR could classify DNA structural states based not only on protein-induced changes but also on environmental variations introduced by the DNA itself, we trained the model on two DNA constructs labeled with Cy3 at the 5’ end: a 32-nt (nucleotide) gap DNA (serving as a DNA synthesis template) and a nicked DNA (a product of DNA synthesis) (Figure 4a). This setup enabled us to investigate whether the structural transition from a gap to a nick influences the local fluorophore environment in the absence of additional binding proteins. We suspected that the resulting environmental change could provide the source of smDeepFLUOR to fluorescence signals

**Figure 4.**
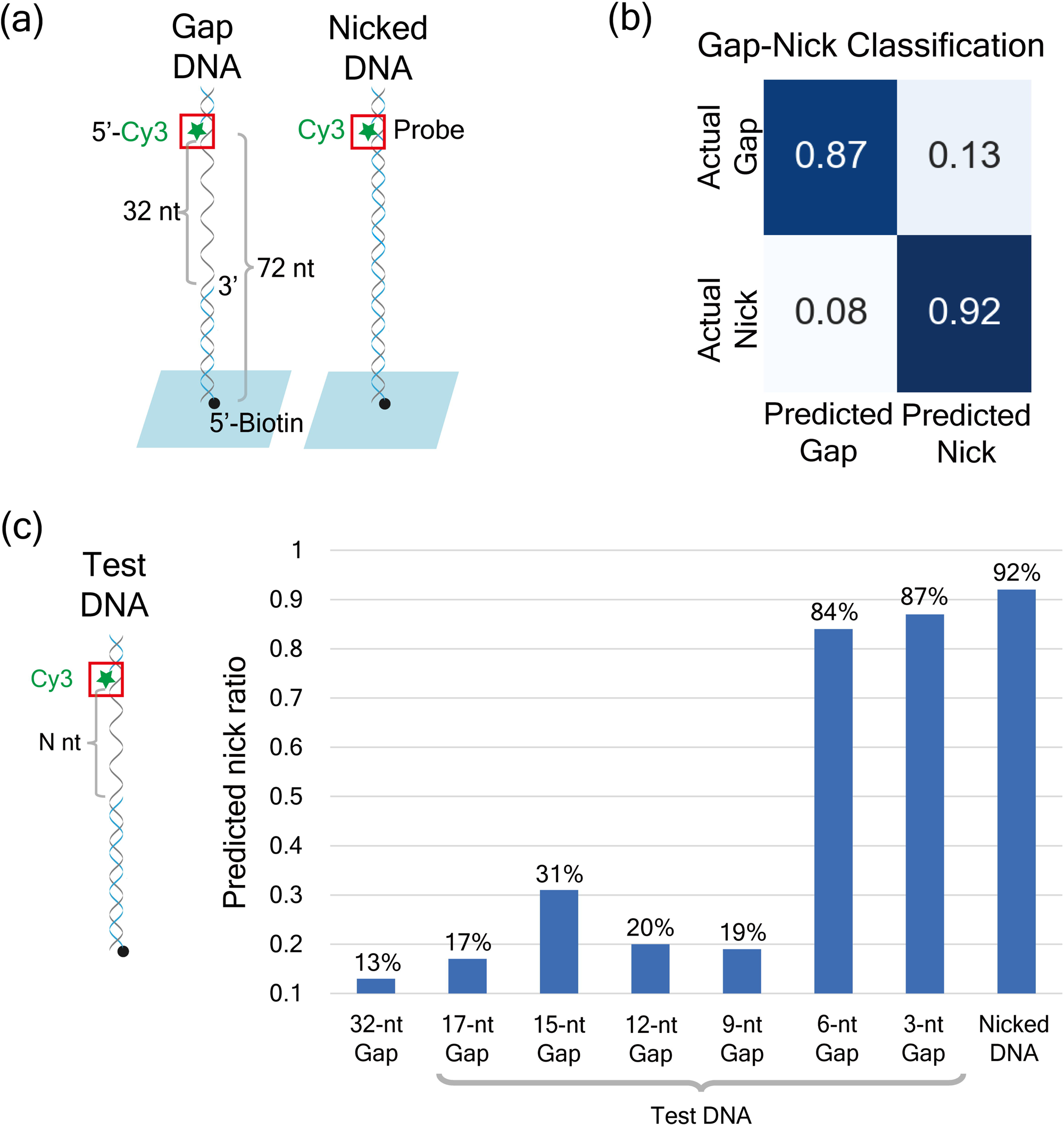
Classification of two different structure of DNAs, gap DNA and nicked DNA. (a) Schematics of Cy3-labeled 32-nt gap DNA and nicked DNA. Fluorescence signals from Cy3 were used as input for model training (green star in red box). (b) In the absence of PCNA, RFC, and dNTPs, smDeepFLUOR achieved 87% accuracy in classifying gap DNA and 92% accuracy for nicked DNA. (c) DNA constructs for various gap lengths (N = 17 nt, 15 nt, 12 nt, 9 nt, 6 nt, and 3 nt). Prediction results for these test DNAs were obtained using the model trained on 32-nt gap DNA and nicked DNA. The 17-nt, 15-nt, 12-nt, and 9-nt gap DNAs were predicted as nicked DNA with accuracies of 17%, 31%, 20%, and 19%, respectively. In contrast, the misclassification rate increased sharply for shorter gaps: the 6-nt gap and 3-nt gap DNAs were predicted as nicked DNA with 84% and 87% accuracy, respectively.

Using Cy3 intensity profiles obtained via TIRF microscopy, smDeepFLUOR was trained on Cy3 labeled 32-nt gap DNA and nicked DNA. It achieved 87% accuracy in identifying the gap DNA and 92% accuracy for the nicked DNA, resulting in an overall classification accuracy of 89% in distinguishing between the two structural states (Figure 4b). To evaluate whether the model could generalize to mixed populations, the model was tested on 1:1 and 9:1 mixtures of the two DNA constructs. The predicted distributions closely matched the expected ratios of 45:55 and 92:8, respectively (Figure S2). These results suggest that smDeepFLUOR can reliably detect structural transitions between 32-nt gap DNA and nicked DNA with minimal labeling.

Importantly, we applied the model, trained solely on 32-nt gap and nicked DNA, to DNA molecules containing shorter gaps of 17, 15, 12, 9, 6, and 3 nt. The 17-, 15-, 12-, and 9-nt gap constructs were consistently classified as 32-nt gap DNA with accuracies ranging from 69% to 83% (Figure 4c). In contrast, the 6-nt and 3-nt gap DNA molecules were predominantly predicted as nicked DNA, achieving 84% and 87% accuracy, respectively (Figure 4c). Interestingly, these findings align with previous reports of NAIFE occurring within a 10-nt range: short single-stranded segments can measurably alter the fluorophore’s local environment, even in the absence of proteins^5^. While the change in fluorescence intensity in our DNA structures may be too subtle to be detected by conventional methods, it is likely that smDeepFLUOR can recognize such variations through its sensitivity to spatiotemporal emission patters.

### Classification of the DNA synthesis by distinct DNA polymerases

To assess smDeepFLUOR’s ability to detect the transition of gap DNA to nicked DNA by DNA polymerase, we first trained the model using gap and nicked DNA molecules in the presence of proliferating cell nuclear antigen (PCNA), replication factor C (RFC), and dNTPs, key components required for DNA synthesis, but in the absence of DNA polymerase. The classification accuracies under these conditions were slightly reduced, with 81% for gap DNA and 82% for nicked DNA (Figure 5a), suggesting that DNA synthesis components (PCNA, RFC, and dNTPs) other than DNA polymerase have minimal influence on smDeepFLUOR classification performance.

**Figure 5.**
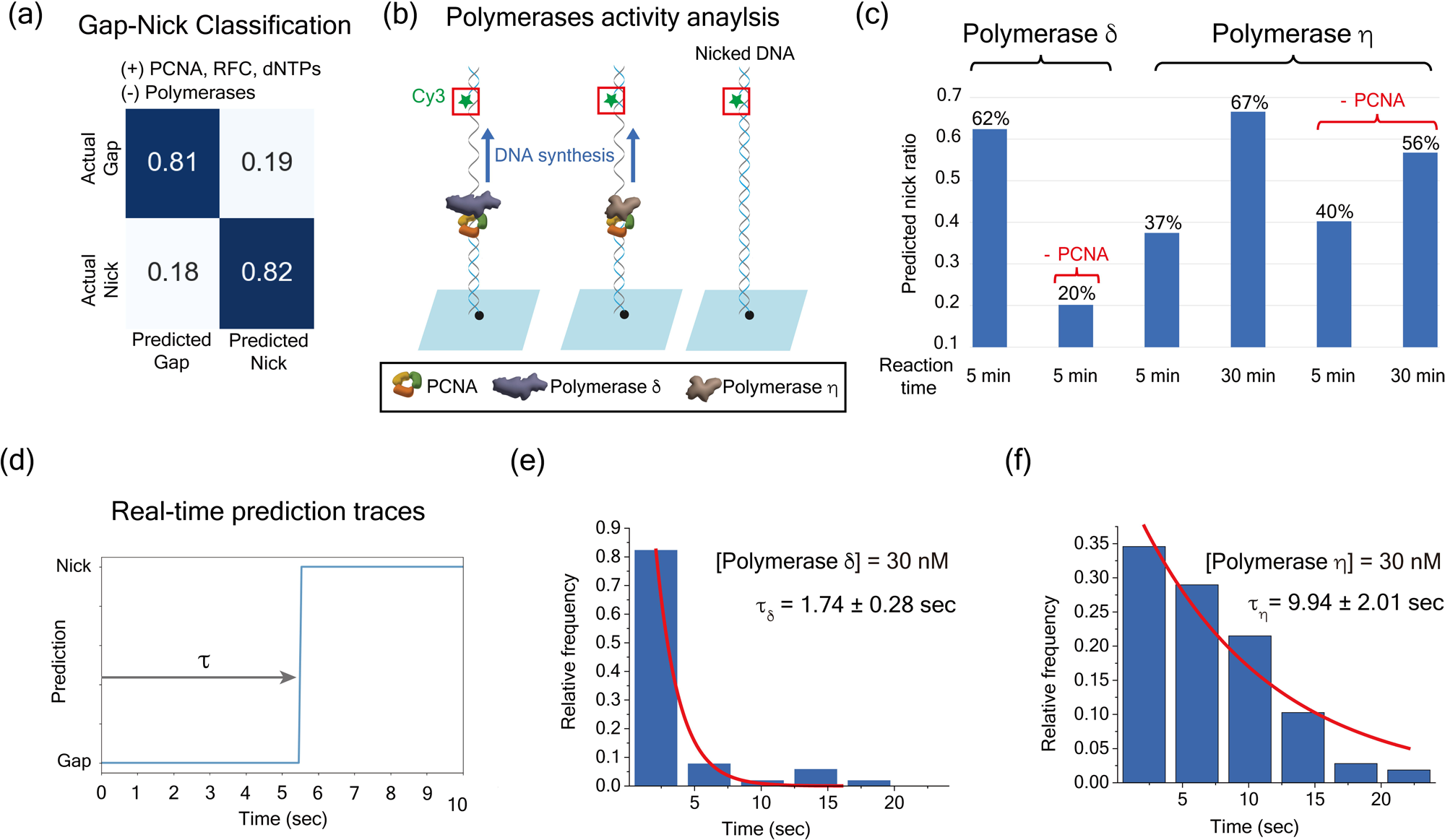
DNA synthesis by DNA polymerases monitored with smDeepFLUOR. (a) In the presence of PCNA, RFC, and dNTPs, smDeepFLUOR achieved classification accuracies of 81% for gap DNA and 82% for nicked DNA. (b) Schematic of the polymerase activity assay illustrating the gap-to-nick transition of DNA in the presence of DNA polymerases. (c) Predicted nick-to-gap DNA ratios under various conditions. The smDeepFLUOR predicted 62% nick formation by DNA polymerase δ at 5 min, and 37% and 67% by DNA polymerase η at 5 and 30 min, respectively. Without PCNA, predicted nick ratios were 20% for DNA polymerase δ at 5 min, and 40% and 56% for DNA polymerase η at 5 and 30 min, respectively. (d) Representative real-time trace of smDeepFLUOR predictions showing the measured gap-to-nick transition time (τ), including the association time of DNA polymerases. (e, f). The mean gap-to-nick transition time was 1.7 ± 0.3 seconds for DNA polymerase δ and 9.9 ± 2.0 seconds for DNA polymerase η, as shown in the resulting histograms.

We then applied smDeepFLUOR to analyze the activities of two distinct human DNA polymerases: polymerase δ^20^ and polymerase η^21^ (Figure 5b). DNA polymerase δ exhibits a stronger PCNA-dependent association and a higher DNA synthesis rate than polymerase η, whose activity is relatively independent of PCNA^22^. Despite identical average intensity of the probe Cy3 on DNA (Figure S3), smDeepFLUOR successfully detected their distinct biochemical behaviors in the classification of gap and nicked DNA.

In the presence of all components, including polymerase δ, the model predicted that 62% of gap DNA transitioned to nicked DNA within 5 min after reaction initiation. In contrast, in the absence of PCNA, only 20% of gap DNA was classified as nicked DNA (Figure 5c). As expected, smDeepFLUOR also detected the gap-to-nick transition by DNA polymerase η, albeit with a sixfold slower rate than polymerase δ at 30 min reaction time (Figure 5c). Moreover, the smDeepFLUOR classification indicated that polymerase η-mediated DNA synthesis occurred largely independent of PCNA, as the predicted nick ratios were nearly identical regardless of PCNA presence at the same reaction time (Figure 5c). The results demonstrate that smDeepFLUOR accurately classifies gap and nicked DNA, in strong agreement with previously reported biochemical data^22, 23, 24^.

Furthermore, real-time prediction trajectories revealed that the nick transition time for gap (Figure 5d; Figure S4) after 30 nM polymerase δ injection (τ_δ_ = 1.74 ± 0.28 sec; Figure 5e) was significantly faster than that of 30 nM polymerase η (τ_η_ = 9.94 ± 2.01 sec; Figure 5f). The nick transition time (τ) reflects both the association and synthesis phases, but it is primarily governed by the association step, as the synthesis rates, 100 s^-1^ for polymerase δ and 67 s^-1^ for polymerase η^22^, are rapid for a 32-nt synthesis. Our observation is consistent with previous reports^22, 23, 24^ indicating that the association rate of polymerase η to PCNA-loaded DNA is generally slower and weaker than that of DNA polymerase δ. Together, these results suggest that smDeepFLUOR can serve as a highly sensitive and quantitative tool for monitoring DNA synthesis or excision in a minimally labeled system containing only a single fluorophore on the template strand.

## Discussion

Single-molecule fluorescence imaging offers deep insights into biomolecular interactions. However, its interpretability has long been constrained by the limited dimensionality of fluorescence signals, which are typically confined to fluorescence intensity or lifetime. In this study, we present smDeepFLUOR, a deep learning– based classification framework designed to extract high-content spatiotemporal features from the fluorescence emission dynamics of a single fluorophore, enabling discrimination beyond the capabilities of conventional intensity-based analyses. Remarkably, smDeepFLUOR maintains robust performance without the need for engineered features or multiplexed labeling, even when the mean fluorescence intensities between distinct molecular states were indistinguishable by conventional methods. This underscores the existence of subtle but learnable information embedded within the raw fluorescence dynamics.

Recent deep learning approaches in single-molecule imaging have primarily focused on exploiting dynamic photophysical properties of fluorophores, such as blinking kinetics^13^, or on employing engineered and structured point spread functions (PSFs) that encode dipole orientation or wavefront distortions^7, 8, 25^. In contrast, our work demonstrates that meaningful classification of biomolecular states can be achieved from subtle, intrinsic variations in the PSF arising from changes in the local environment of a single fluorophore. To our knowledge, this is the first demonstration of PSF-mediated state classification without any explicit optical engineering or photophysical modulation.

To elucidate the origin of this discriminatory signal captured by smDeepFLUOR, we conducted control experiments using Cy3-labeled double-stranded DNA (dsDNA) of varying total lengths, while maintaining the local fluorophore environment identical (Figure S5). In these conditions, smDeepFLUOR failed to differentiate among the dsDNA molecules, which indicates that global structural properties or molecule-wide diffusion are unlikely to contribute to classification. Instead, experiments with variable gap lengths revealed that classification accuracy showed a sharp distinction between 9-nt and 6-nt gaps, suggesting that the fluorophore is responsive to localized environmental differences within a spatially restricted range (Figure 4c). This is reminiscent of distance-dependent effects observed in mechanisms such as PIFE and NAIFE, which typically operate over distances of ∼4 nm^26^ or ∼7–9 nucleotides^5^. However, smDeepFLUOR achieved this discrimination without relying on abrupt intensity change, implying that it is likely to detect subtle, spatially confined variations in the emission profile rather than apparent intensity shifts.

Future extensions of smDeepFLUOR may include incorporating transfer learning^27, 28^ or self-supervised training^29^ schemes to adapt the model to new molecular targets with limited annotation data. In particular, unsupervised classification of the spatiotemporal patterns of fluorescent molecule PSFs is of great significance. It has been observed that even slight changes in the local environment can induce detectable shifts in the emission profile of a single fluorophore, giving rise to distinct spatiotemporal PSFs. Therefore, if we can develop analysis methods that do not require training data, such as interpreting pixel-wise temporal fluctuations through autocorrelation or cross-correlation matrices^30^, it may become feasible to identify arbitrary and previously undefined molecular states. In addition to analytical approaches based on interpretable mathematical functions, the application of self-supervised training methods, such as masked autoencoders (MAE)^31^ in a label-free manner, could offer a powerful framework for uncovering latent molecular states previously inaccessible by conventional means.

In conclusion, our findings establish smDeepFLUOR as a robust analytical framework capable of decoding hidden molecular information from seemingly homogeneous single-fluorophore signals. Its ability to operate with minimal labeling, generalize across independent experimental batches, and track single-molecule dynamics in real time makes smDeepFLUOR a valuable tool for probing subtle molecular interactions in diverse biological systems. Future work may integrate explainability techniques (e.g., saliency maps^32^ or attention mechanisms^33^) to further clarify which aspects of the fluorescence signal drive classification, potentially revealing new physical insights. Furthermore, extending smDeepFLUOR to broader classes of biomolecules, time scales, and imaging platforms may open new frontiers in real-time, minimally invasive single-molecule analysis.

## Methods

### Quartz preparation

The quartz slides were cleaned through sequential sonication following a previously reported protocol^34^. The cleaning steps included in 10% Alconox solution for 20 minutes, followed by deionized water (Milli-Q) for 5 minutes, acetone for 15 minutes, KOH for 20 minutes, and methanol for 10 minutes. They were then flame-treated with a torch for 30 seconds. After cleaning, the slides were incubated in a 2% (v/v) solution of (3-aminopropyl) triethoxysilane (Sigma) for 20 minutes. A mixture of polyethylene glycol succinimidyl valerate (PEG-SVA, MW 5000, Laysan Bio) and Biotin-PEG-SVA (MW 5000, Laysan Bio) at a mass ratio of 40:1 was prepared in 0.1 M sodium bicarbonate buffer (pH 8.3). This solution was applied to the quartz slide surface and incubated for 6 hours. Finally, the functionalized slides were rinsed with deionized water and dried with nitrogen gas.

### Total internal reflection fluorescence (TIRF) microscopy

All the single-molecule TIRF images in this study were acquired with the 1.6X magnifier in a prism-type TIRF microscope (Olympus IX-71, water-immersion 60X objective NA = 1.2), EMCCD (ImagEM C9100-13, Hamamatsu), MetaMorph 7.6 (Molecular Devices) imaging software and a Dual View optical setup (DV2, Photometrics) if required. Fluorophores were excited using the laser lines (Cobolt MLD 488 nm-300 mW for mNG, Cobolt DPL 532 nm-100 mW for Cy3, and CrystaLaser 638 nm-70 mW for Cy5 and AF647). The fluorescence signal was filtered with each emission filters (Chroma, ET525/50 m; Semrock, 585/40; Semrock, FF01-676/37-25).

### Single-molecule experiment conditions

#### Poly(A) binding protein (PABP) and DNA/RNA hybrids

We prepared the mNG-PABP, non-labeled PABP, and RNA/DNA hybrid templates following the protocol described in a previous study^35^. Poly(A)_25_, containing a 25-nucleotide poly(A) tail was used for nonspecific classification, while Poly(A)_60_, with a 60-nucleotide poly(A) tail, was used for protein-binding classification. After PEG-biotinylated flow chamber preparation, streptavidin (0.05 mg/ml, Sigma-Aldrich) in blocking buffer (20 mM Tri-HCl, pH 7.5, 2 mM EDTA, 50 mM NaCl, and 0.0025% Tween 20(v/v), all from Sigma) was introduced into the chamber and incubated for 5 min. Following free streptavidin removal by washing with blocking buffer, Cy5-conjugated RNA/DNA hybrid (10 pM) in blocking buffer was injected into the channel and incubated for 10 min before being washed with reaction buffer (25 mM Tris-HCl pH 7.5, 100 mM l-glutamic acid potassium salt monohydrate, 5 mM MgCl_2_, 0.0025% [v/v] Tween 20, 0.01% Triton X-100, all from Sigma). Next, mNG-PABP (200 pM) or non-labeled PABP (10 nM) in reaction buffer was introduced and incubated for 5 min. To acquire images of nonspecific bound PABP, mNG-PABP (1 nM) was incubated in the chamber for 30 min in the absence of RNA/DNA hybrid. Finally, fluorescence images were acquired in the presence of an oxygen scavenger system [2 mM Trolox (Sigma), 5 mM PCA (Sigma), and 200 nM rPCO (Oriental Yeast)].

#### Human Pol **η** purification

Human full-length DNA polymerase η (PolH) was expressed in E. coli BL21 (DE3) cells using an N-terminal (His)\₆-SUMO tag from the pE-SUMO-pro expression vector (LifeSensors). Cells were cultured in 2YT medium at 24°C to an OD_600_ of 0.8, induced with 0.1 mM isopropyl β-D-1-thiogalactopyranoside (IPTG), and incubated for an additional 19 hours at 16°C. Cells were harvested by centrifugation and resuspended in lysis buffer [50 mM Tris-HCl (pH 8.0), 750 mM NaCl, 40 mM imidazole, 5 mM β-mercaptoethanol, 0.2% NP-40, 1 mM PMSF, 5% glycerol, and 1 EDTA-free protease inhibitor tablet per 50 mL (Roche, UK)]. All subsequent steps were performed at 4°C. Cells were lysed enzymatically with 2 mg/mL lysozyme and mechanically by sonication. The lysate was clarified by centrifugation (22,040 ×Dg, 60 min), and the supernatant was loaded onto a 5 mL HisTrap HP affinity column (Cytiva) pre-equilibrated with buffer A [50 mM Tris-HCl (pH 7.5), 500 mM NaCl, 40 mM imidazole, 5 mM β-mercaptoethanol, and 5% glycerol]. The column was washed with 50 mL of buffer A and eluted using a linear gradient of buffer B [same as buffer A, but with 500 mM imidazole]. PolH eluted at approximately 310 mM imidazole. Peak fractions were pooled and dialyzed overnight in dialysis buffer [50 mM Tris-HCl (pH 7.5), 500 mM NaCl, 5 mM β-mercaptoethanol, and 5% glycerol] in the presence of SUMO protease (LifeSensors) to remove the SUMO tag and yield native PolH. The dialyzed sample was re-applied to the HisTrap column, and the untagged PolH was collected from the flow-through.

PolH-containing fractions were pooled, concentrated, and further purified by size-exclusion chromatography using a HiLoad 16/600 Superdex 200 pg column (Cytiva) equilibrated with storage buffer [50 mM Tris-HCl (pH 7.5), 300 mM NaCl, 10% glycerol, and 1 mM DTT]. Final protein fractions were concentrated, flash-frozen in liquid nitrogen, and stored at −80°C.

#### Human Pol **δ** purification

Polδ WT were expressed in Sf9 insect cells and purified as previously described^20^. Briefly, Human Polδ was expressed in Sf9 insect cells using a baculovirus system. Bacmid DNA encoding all four subunits was transfected into Sf9 cells to generate viral stocks, which were amplified up to P3 virus stock. Cells were transfected with P3 virus and harvested after 72 hours. Cells were lysed and centrifuged and the clear lysate was loaded on to HisTrap affinity column. Eluted proteins were further purified via anion exchange (Mono Q) and size-exclusion chromatography (Superdex 16/600 200 pg). Fractions containing all Polδ subunits were pooled, concentrated, flash-frozen, and stored at −80°C.

#### PCNA purification

N-terminally His-tagged PCNA WT was expressed in E. coli and purified as previously described^36, 37^. Briefly, PCNA plasmid was expressed in BL21 (DE3) Escherichia coli cells. The protein was purified using a HisTrap HP affinity column, followed by ion exchange chromatography on a HiTrap Q column, and finally by size-exclusion chromatography using a HiLoad 16/600 Superdex 200 pg column. Purified protein fractions were pooled, flash-frozen, and stored at −80°C.

#### RFC purification

ΔN-RFC (amino acids: 554−1148 in RFC1) were expressed in E. coli and purified as previously described^20^. Briefly, ΔNRFC all three plasmids containing all five subunits was co-expressed in E. coli BL21(DE3). Cells were grown in 8DL of TB medium at 25°C, induced with 0.2 mM IPTG at OD_600_ of 0.8, and incubated for 24 hours at 16°C. After harvesting, cells were lysed and the clarified lysate was purified using HisTrap affinity chromatography, followed by StrepTrap chromatography to remove improperly assembled complexes. The flow-through containing the correctly assembled ΔNRFC complex was further purified by heparin affinity and size-exclusion chromatography. Fractions with right stoichiometric ΔNRFC were pooled, concentrated, and flash-frozen, and stored at −80°C.

#### DNA polymerases with gap and nicked DNA

DNA oligos were purchased from IDT (Coralville, USA; Table S1). To immobilize DNA templates on the chamber surface, we conducted following the same protocol as the previous PABP binding experiments, with one modification: after streptavidin incubation, Cy3-conjugated gap DNA or nicked DNA (1 pM) in blocking buffer was injected into the channel. Fluorescently labeled molecules were incubated in the chamber for 10 min before being washed with reaction buffer. For the polymerase activity experiments, reaction buffer containing 1 mM ATP, 10 μM ADP, 1 mM DTT, 100 μg/ml BSA, 100 μM each dNTP, 5 nM PCNA, 2.5 nM RFC, and either 30 nM polymerase δ or 90 nM polymerase η were introduced into the chamber. Control experiments were performed in identical conditions, excluding specific interacting molecules as detailed in the main text. Fluorescence images were acquired using in the same imaging buffer containing oxygen scavenger system.

### Single particle cropping from TIRF microscopy images

Single particle detection and cropping were performed using imageJ plugin mosaic^38^. Prior to running the plugin, background intensity was uniformly adjusted using rolling ball background subtraction in imageJ. Particle detection was conducted with the following hyperparameters: radius = 3, cutoff = 0.001, and absolute threshold = 250, 300, 350, with smear particles below the threshold being excluded. For particle tracking, we applied displacement = 1 and 2.5 and link range = 1. Except for the determination of colocalization between particles, the displacement hyperparameter was set to 1, meaning that a particle was allowed to move a maximum of one pixel between consecutive frames. Since smDeepFLUOR predicts states based on 10-frame sequence, traces shorter than 10 frames were discarded.

### Image Standardization

Standardization of a data is crucial for increasing training performance by ensuring faster and more stable convergence during gradient descent, improving optimization, and enabling more effective weight initialization. It also boosts the performance of activation functions like ReLU which we used in our model, enhances the effectiveness of batch normalization, and helps reduce overfitting by ensuring fair contribution from all features. and efficiency of gradient descent. Intensity from each pixel was standardized within a 7 x 7 x 10 cube (490 pixels). The equation used for the standardization is as follows:

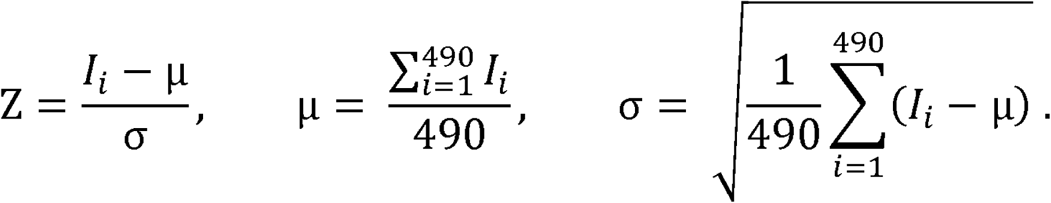

Here, *l_i_* is the intensity of each pixel and *Z* is the z-score of *l_i_* in each distribution with mean μ and standard deviation σ.

### Training and validation

To construct the full dataset, particle images (7 x 7 x N, where N represents the number of frames before fluorescence bleaching) were cropped from spots that were tracked for more than 10 frames in a field of view for each state. For each classification task, the full dataset was randomly split into an 80:20 ratio to create the training and test sets. During classification training, the model was trained using 7 x 7 x 10 cubes sequentially extracted from the training set, while validation was performed using 7 x 7 x 10 cubes extracted from the test set. If the training data exceeded 50,000 cubes of 7 x 7 x 10, a random subset of 50,000 cubes was selected in training set.

For prediction results not shown in real-time, the final prediction was determined by averaging the predictions of 7 x 7 x 10 cubes extracted from 7 x 7 x N images of a single particle. These results were used to construct all confusion matrices. The statistics of real-time predictions, before averaging whole 7 x 7 x N images, were summarized for all experiments (Figure S6). In single-molecule predictions, all prediction results from the same particle were utilized to determine their states. In contrast, single-molecule real-time prediction data were averaged over 10 successive 7 x 7 x 10 cubes to capture state transitions. While real-time prediction does not entirely filter outliers or fluctuations within a trace, the overall trend of the predictions remained consistent. Even when fluctuations occurred within specific 7 x 7 x 10 cubes from 7 x 7 x N images of the same particle, the predictions were generally stable and reliable.

The overall model architecture remained consistent across all classification tasks (Table S2), though slight variations in dropout ratios were applied for different tasks. During training, various dropout ratio ranges were tested, and the optimal value for each specific task was selected. The total number of training and test samples used for all classification task is summarized in Table S3. Model training was performed using Google Colab Pro with an A100 GPU. Scikit-learn^39^ and tensorflow^40^ were used for data processing and model building.

Lastly, to confirm that smDeepFLUOR specifically learns fluorescence-related features, background images were tested during training. 7 x 7 x 10 background images adjacent to original fluorescence signals were collected to determine whether classification was independent of background information. When attempting to classify background images from different experimental conditions using the same smDeepFLUOR architecture, the model performed poorly, demonstrating that smDeepFLUOR detects intrinsic fluorescence-dependent differences rather than background variations (Figure S7).

## Data Availability

The data shown in this article are available from the corresponding authors upon a request.

## Code Availability

The python codes used for analysis in this article are available from the corresponding authors upon a request.

## Acknowledgments

This work was supported by the National Research Foundation of Korea funded by the Ministry of Science and ICT (RS-2023-00280169 and RS-2023-00218927) and by Basic Science Research Institute Fund, whose NRF grant number is RS-2021-NR060139.

## Author Contributions

Conceptualisation: J.L. and J.B.L. Methodology: J.L. Single-molecule experiments: J.L., B.K., G.B. Protein purification: B.K., M.T., V.S.R. Writing: J.L. and J.B.L. Editing: J.L., B.K., S.M.H. and J.B.L.

## Notes

### Competing Interest Statement

The authors have declared no competing interest.

