## Supplementary information for "smDeepFLUOR: Single-Molecule Deep Learning Fluorescence Classification"

**Tables**

- Table S1. Oligos for DNA polymerase activity analysis experiment.
- Table S2. Detailed architecture of smDeepFLUOR.
- Table S3. Detailed data on training sets and test sets for each classification task.
- Table S4. Oligos for supplementary figure 5, Cy5-labeled dsDNA length classification.

**Supplementary Figures**

- Figure S1. PABP binding prediction.
- Figure S2. Gap-Nick classification.
- Figure S3. Fluorescence intensity of Cy3-gap DNA and nicked DNA.
- Figure S4. Analysis of real-time Gap - Nick prediction.
- Figure S5. Training results of various Cy5-labeled double-stranded DNAs (dsDNA)
- Figure S6. Statistics of single-molecule predictions.
- Figure S7. Training results of background noise images.

**Table S1. Oligos for DNA polymerase activity analysis experiment.** All oligos were purchased from IDT (Integrated DNA technology).

| **Name** | **Sequence and modification** |
| --- | --- |
| Cy3 oligo | /5Cy3/ GGT CGT CAG TGC TGG GCG TC /3Dig_N/ |
| Biotin oligo (32nt gap) | /5Biosg/ GTG CAT AAG CAG TCT TAG TGG TCT GAT GTA TGG ACT TAC C |
| Biotin oligo (17nt gap) | /5Biosg/ GTG CAT AAG CAG TCT TAG TGG TCT GAT GTA TGG ACT TAC CTA TCA CGG ATT CTG A |
| Biotin oligo (15nt gap) | /5Biosg/ GTG CAT AAG CAG TCT TAG TGG TCT GAT GTA TGG ACT TAC CTA TCA CGG ATT CTG AAG |
| Biotin oligo (12nt gap) | /5Biosg/ GTG CAT AAG CAG TCT TAG TGG TCT GAT GTA TGG ACT TAC CTA TCA CGG ATT CTG AAG AGA |
| Biotin oligo (9nt gap) | /5Biosg/ GTG CAT AAG CAG TCT TAG TGG TCT GAT GTA TGG ACT TAC CTA TCA CGG ATT CTG AAG AGA GGG |
| Biotin oligo (6nt gap) | /5Biosg/ GTG CAT AAG CAG TCT TAG TGG TCT GAT GTA TGG ACT TAC CTA TCA CGG ATT CTG AAG AGA GGG ATG |
| Biotin oligo (3nt gap) | /5Biosg/ GTG CAT AAG CAG TCT TAG TGG TCT GAT GTA TGG ACT TAC CTA TCA CGG ATT CTG AAG AGA GGG ATG AGG |
| Complementary oligo | GAC GCC CAG CAC TGA CGA CCG TGC CTC ATC CCT CTC TTC AGA ATC CGT GAT AGG TAA GTC CAT ACA TCA GAC CAC TAA GAC TGC TTA TGC AC |

**Table S2. Detailed architecture of smDeepFLUOR.**

| **Layer (type)** | **Layer settings** | **# Param** |
| --- | --- | --- |
| Conv3d | Input shape (7,7,10,1), number of filters: 32, kernel size: (2,2,1), activation='relu' | 160 |
| Batch Normalization |  | 128 |
| Dropout | Dropout rate: 0.4, 0.6 | 0 |
| Conv3d | Input shape (6,6,10,32), number of filters: 16, kernel size: (3,2,1), activation='relu' | 3088 |
| Batch Normalization |  | 64 |
| Dropout | Dropout rate: 0.2, 0.4 | 0 |
| Conv3d | Input shape (4,5,10,16), number of filters: 16, kernel size: (2,2,10), activation='relu' | 20512 |
| Batch Normalization |  | 128 |
| Dropout | Dropout rate: 0.4, 0.6 | 0 |
| Flatten | Output shape: (none, 384) | 0 |
| Dense | Units: 16, activation='relu', regularizer=l2 | 6160 |
| Dense | Units: 2, activation='softmax' | 34 |

**Table S3. Detailed data on training sets and test sets for each classification task.** Summary of the total number of training and test datasets (7x7x10, tensors).

| **Images** | **# data for training** | **# data for test** |
| --- | --- | --- |
| Non-specific mNG-PABP | 40000 | 10293 |
| Specific mNG-PABP | 40000 | 13004 |

| **Images** | **# data for training** | **# data for test** |
| --- | --- | --- |
| DNA/RNA partial duplex Cy5 | 25000 | 10492 |
| DNA/RNA partial duplex Cy5 with mNG-PABP | 17000 | 4285 |
| DNA/RNA partial duplex Cy5 with  Non-labeled PABP | 20000 | 6529 |

| **Images** | **# data for training** | **# data for test** |
| --- | --- | --- |
| Gap DNA Cy3 | 50000 | 57191 |
| Nicked DNA Cy3 | 50000 | 35263 |

**Table S4.** **Oligos for supplementary figure 5, cy5-labeled dsDNA length classification.** All oligos were purchased from Bionics (Seoul, Korea). The 70 bp, 120 bp, and 150 bp dsDNA fragments were generated by PCR amplification from an arbitrary plasmid, whereas the 30 bp dsDNA was prepared by direct annealing of two complementary oligonucleotides.

| **Name** | **Sequence and modification** |
| --- | --- |
| Cy5 oligo | /Cy5/ CCG ATT GCT ACC CTG AAG AAT TTT CCA AA |
| Biotin primer (29bp) | /5Biosg/ TTT GGA AAA TTC TTC AGG GTA GCA ATC GG |
| Biotin primer (70bp) | /5Biosg/ CAA ATT CAT CGC GCG CCC A |
| Biotin primer (120bp) | /5Biosg/ ATC GGT CAG ATA CTG ATT CAC GTT TTC C |
| Biotin primer (145bp) | /5Biosg/ GCA GGG TGC GTT CCA CAA ATT TC |

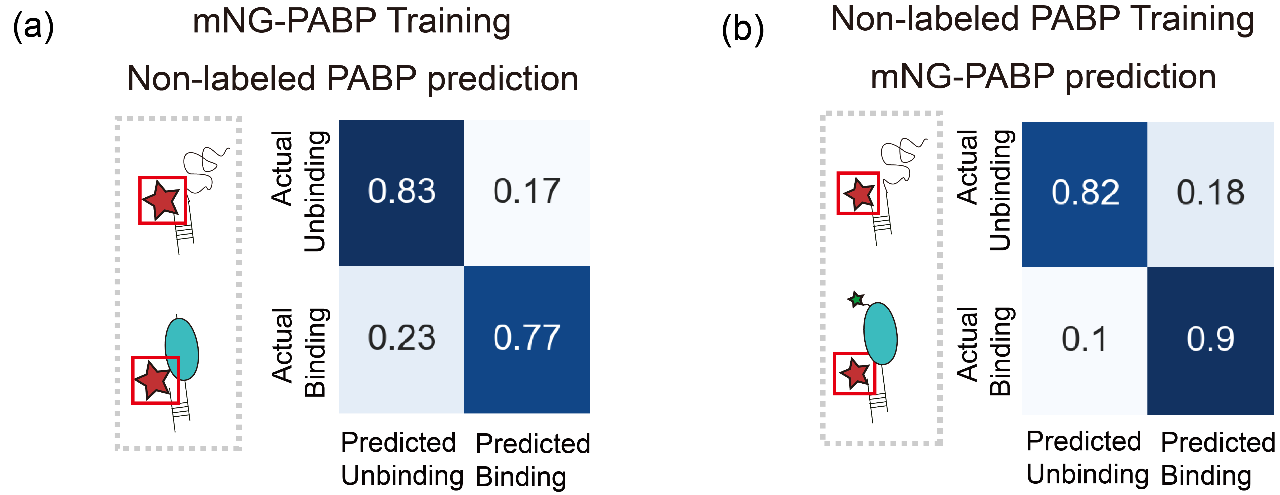

**Figure S1.** **PABP binding prediction**. (a) Prediction results for unlabeled PABP using smDeepFLUOR trained on mNG-PABP. The confusion matrix shows the smDeepFLUOR classified unbound PABP with 83% accuracy and bound PABP with 77% accuracy. (b) Prediction results for mNG-PABP using smDeepFLUOR trained on non-labeled PABP. The smDeepFLUOR achieved 82% accuracy for unbound PABP 90% accuracy for bound PABP.

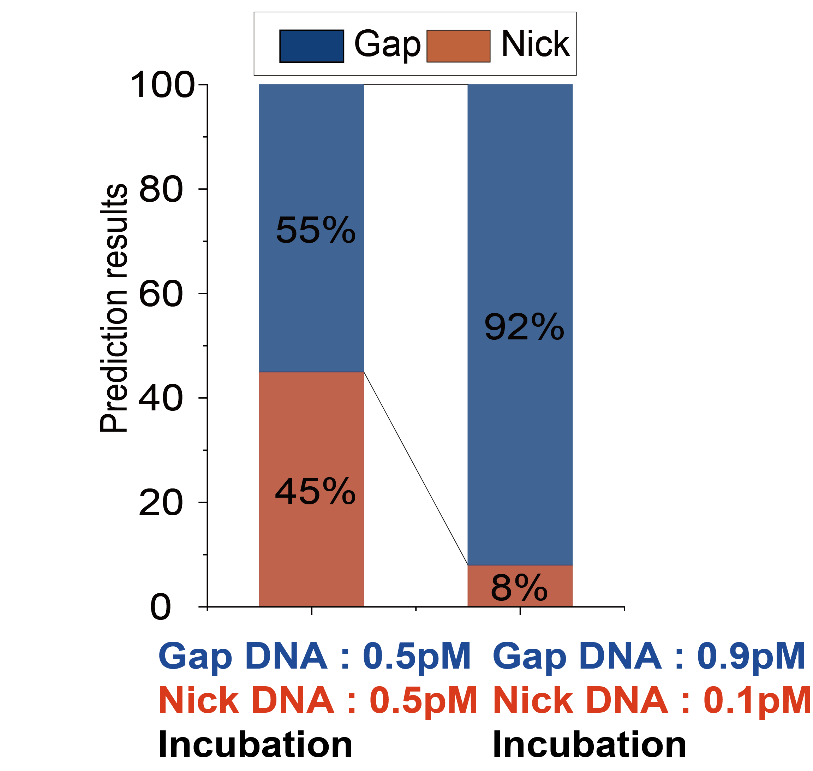

**Figure S2.** **Gap-Nick classification**. Given that smDeepFLUOR classified gap and nick DNA with an average 89% accuracy (Fig. 2b), we tested its performance in mixed populations. When gap and nicked DNA were mixed at 1:1 and 9:1 ratio, smDeepFLUOR identified them with predicted proportions of 45:55 and 92:8, respectively.

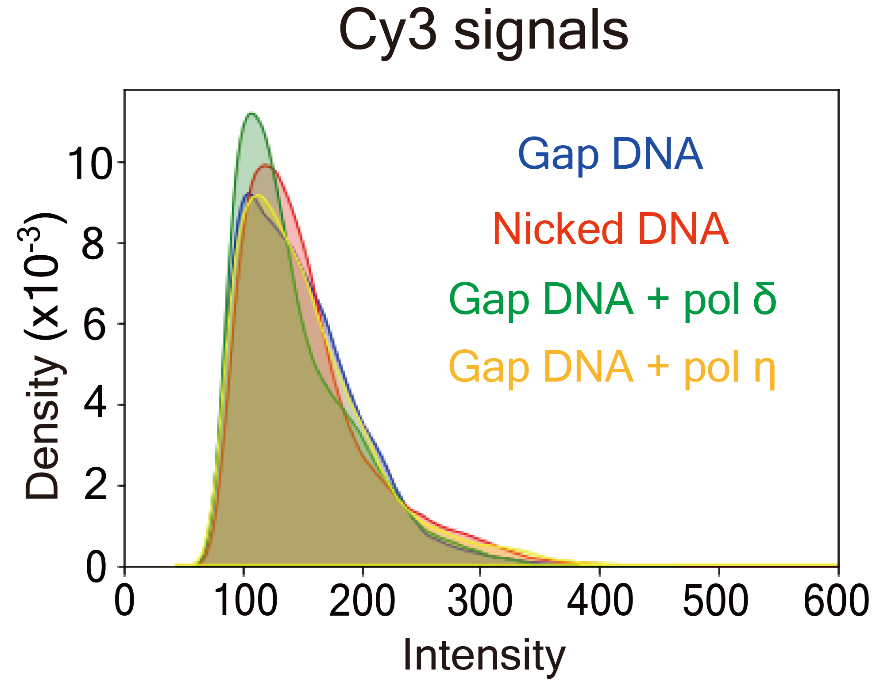

**Figure S3.** **Fluorescence intensity of Cy3-gap DNA and Cy3-nicked DNA**. The average intensity of each particle was calculated by averaging pixel intensities across all frames (7 x 7 x N, where N is the frames before fluorescence bleaching) before normalization. Histograms show Cy3 intensity distributions for Cy3-nicked and gap DNA, with and without polymerases.

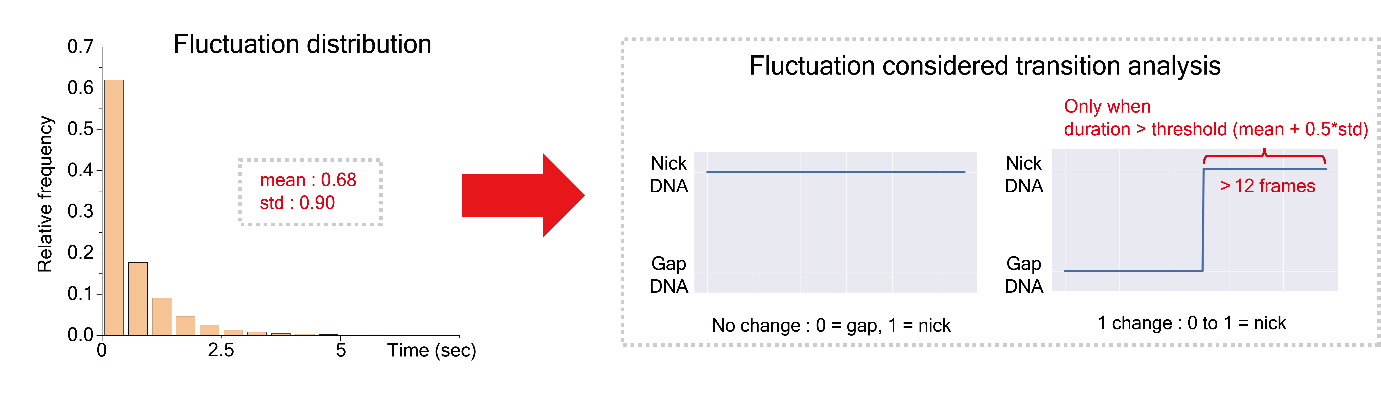

**Figure S4.** **Analysis of real-time gap-nick prediction.** A method for calculating the nick ratio incorporating both population-level predictions and real-time transitions. The histogram (left) summarizes real-time fluctuation statistics from all data, including polymerase δ and polymerase η. A gap-to-nick transition was defined as a true transition only if the post-transition state persisted longer than the mean + 0.5×SD (~12 frames, or 1.2 s); shorter transitions were excluded (right).

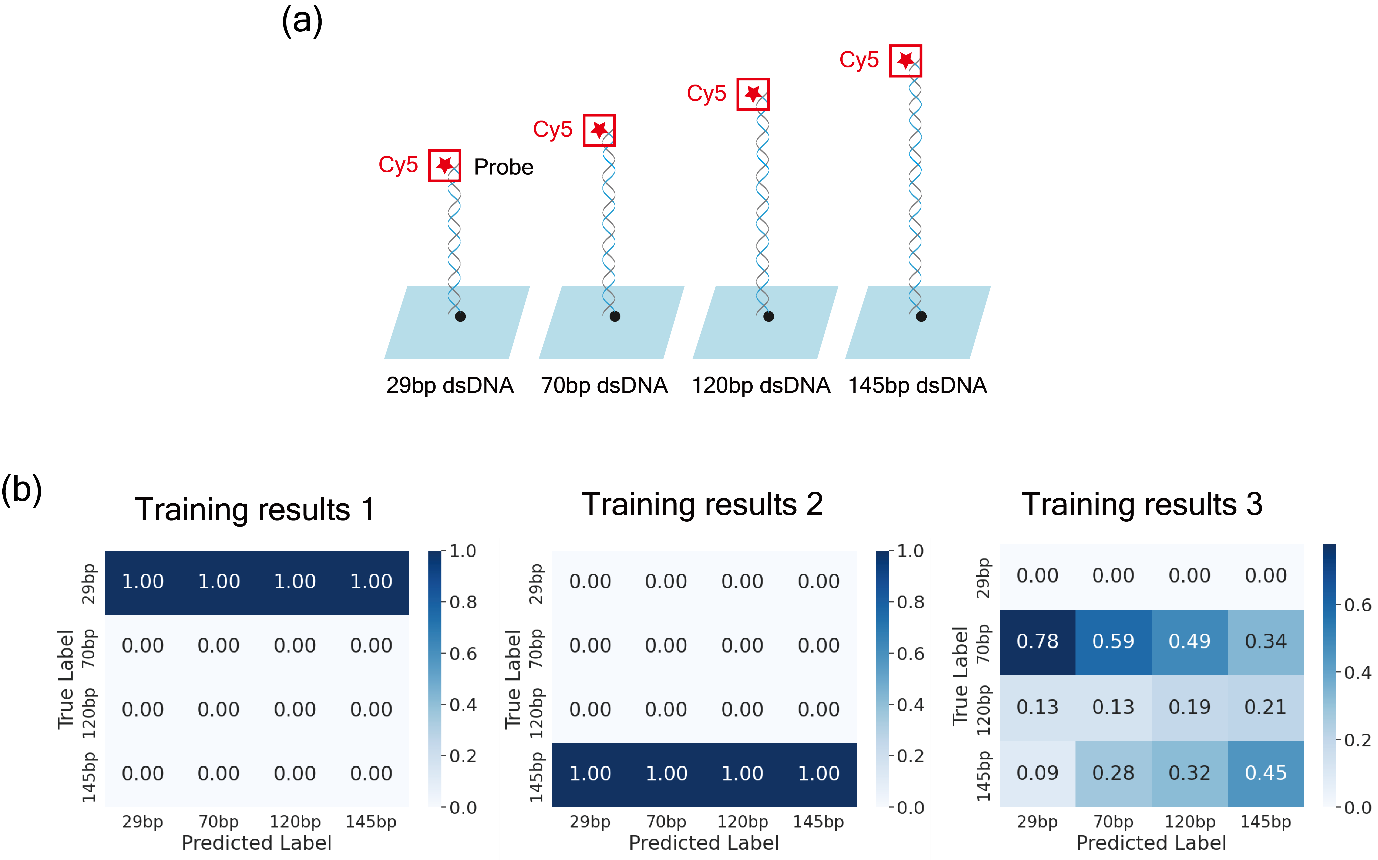

**Figure S5.** **Training results of various Cy5-labeled double-stranded DNAs (dsDNA)**. (a) Schematic illustration of surface-tethered Cy5-labeled double-stranded DNA (dsDNA) constructs of varying lengths (29, 70, 120, and 145 base pairs). Each dsDNA is immobilized on a glass surface and labeled at the top with a Cy5 fluorophore for fluorescence signal acquisition. (b) Confusion matrices from three independent training runs of a model classifying the different dsDNA lengths. Despite using identical model architecture and training configuration, results vary significantly between runs: Training results 1 and 2 show complete prediction collapse to a single class (145 bp and 29 bp, respectively), indicating failure to learn meaningful class distinctions. Training result 3 shows partial class separation but still suffers from significant confusion across all classes, particularly between 70 bp and 145 bp. These results suggest that the dataset lacks sufficient class-separable information for stable supervised learning.

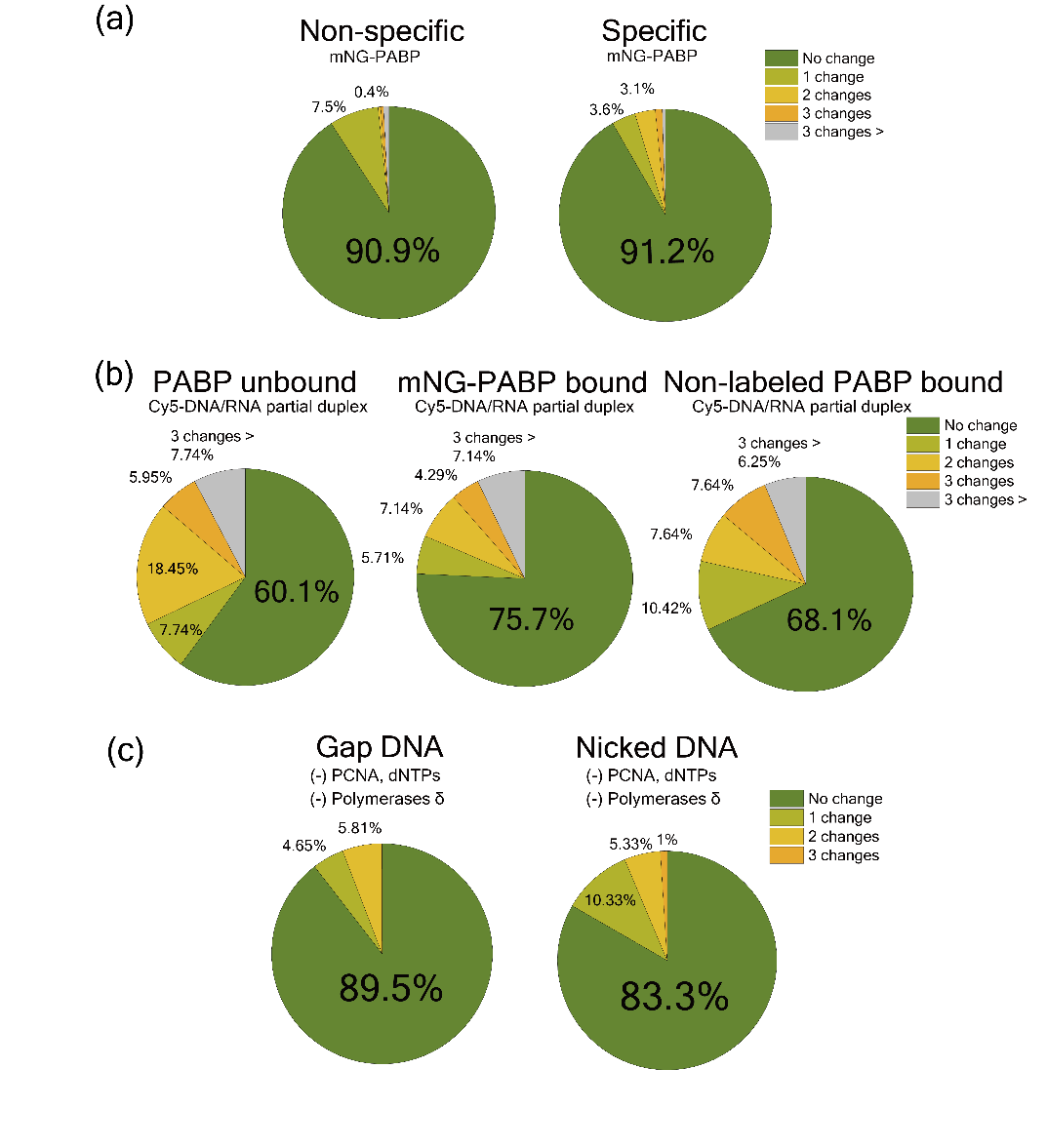

**Figure S6.** **Statistics of single-molecule traces predictions**. The pie charts represent trace-level prediction outcomes from the data used in each confusion matrix, without averaging across individual particles. (a) Prediction fluctuation statistics for mNG traces from nonspecifically bound mNG-PABP (left) and poly(A)-bound mNG-PABP (right). A total of 90.9% and 91.2% of traces, respectively, remained unchanged throughout the analysis. (b) Prediction fluctuation statistics for Cy5 traces in the presence of PCNA and dNTPs. Results are shown for PABP-unbound Cy5-DNA/RNA partial duplex (left), mNG-PABP–bound (middle), and unlabeled PABP–bound duplex (right). The percentages of stable traces were 60.1%, 75.7%, and 68.1%, respectively. (d) Prediction fluctuation statistics for Cy3 traces from gap DNA (left) and nicked DNA (right) in the absence of polymerases. Traces remained unchanged in 89.5% and 83.3% of cases, respectively.

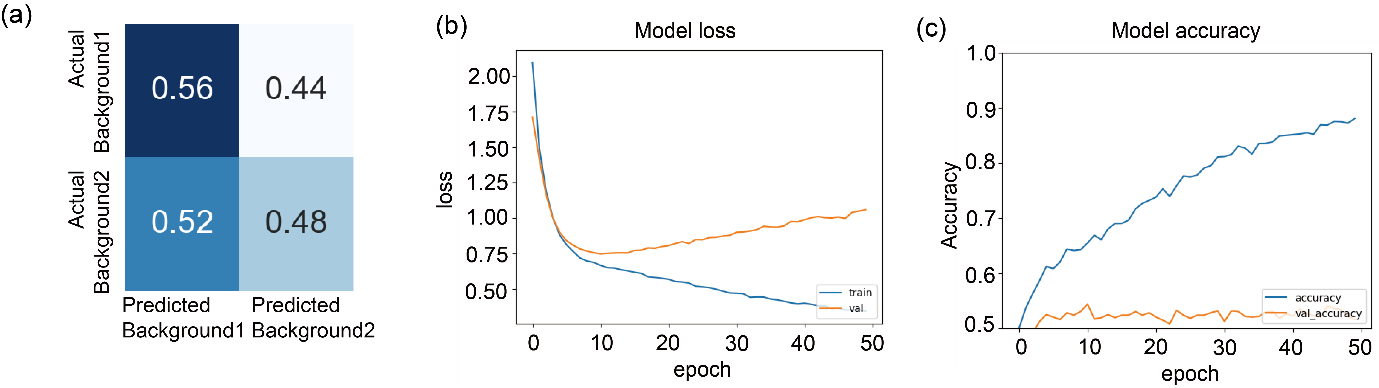

**Figure S7.** **Training results of background noise images**. To assess whether smDeepFLUOR learns different fluorescence states, we cropped random regions for background training. (a) The confusion matrix shows prediction results for background signals from experiments with nonspecifically and specifically bound PABP. The smDeepFLUOR failed to identify any distinguishable pattern from background-only images. (b) Training (blue) and validation (orange) loss over epochs. (c) Training (blue) and validation (orange) accuracy over epochs. When trained on background noise instead of actual fluorescence signals, smDeepFLUOR showed no improvement in validation accuracy or reduction in validation loss, indicating that it does not extract features from background alone.
